# Crystal structures reveal the framework of *cis*-acyltransferase modular polyketide synthases

**DOI:** 10.1101/2023.02.11.528132

**Authors:** Adrian T. Keatinge-Clay, Takeshi Miyazawa, Jie Zhang, Katherine A. Ray, Joshua D. Lutgens, Ramesh Bista, Sally N. Lin

## Abstract

Although the domains of *cis*-acyltransferase (*cis*-AT) modular polyketide synthases (PKS’s) have been understood at atomic resolution for over a decade, the domain-domain interactions responsible for the architectures and activities of these giant molecular assembly lines remain largely uncharacterized. The multimeric structure of the α_6_β_6_ fungal fatty acid synthase (FAS) provides 6 equivalent reaction chambers for its acyl carrier protein (ACP) domains to shuttle carbon building blocks and the growing acyl chain between surrounding, oriented enzymatic domains. The presumed homodimeric oligomerization of *cis*-AT assembly lines is insufficient to provide similar reaction chambers; however, the crystal structure of a ketosynthase (KS)+AT didomain presented here and three already reported show an interaction between the AT domains appropriate for lateral multimerization. This interaction was used to construct a framework for the pikromycin PKS from its KS, AT, and docking domains that contains highly-ordered reaction chambers. Its AT domains also mediate vertical interactions, both with upstream KS domains and downstream docking domains.

## INTRODUCTION

Modular polyketide synthases (PKSs) are among of the largest known proteins^1^. Their tens to hundreds of domains collaborate to synthesize complex polyketides, such as the antibacterials erythromycin and mupiricin, from simple carbon precursors. Whether these domains are organized in a higher-order architecture or as beads on a string has long been debated. Even with the advances in electron microscopy (EM) and protein structure prediction, the complexity of these synthases has impeded their architectural characterization^2-7^.

The two major classes of modular PKSs, *cis*-acyltransferase and *trans*-acyltransferase (*cis*-AT and *trans*-AT) assembly lines, possess similar sets of domains, known as modules (updated definition used throughout)^8-12^. Each module generally adds one monomer to the growing acyl chain and is minimally comprised of an AT domain that selects a carbon building block, an acyl carrier protein (ACP) domain that acquires that building block as well as the growing acyl chain, and a ketosynthase (KS) domain that catalyzes carbon-carbon bond formation between the two. Modules in both classes can also harbor processing domains, such as ketoreductases (KR’s), dehydratases (DH’s), and enoylreductases (ER’s), that modify the most recently added portion of the chain. The AT domains of *cis*-AT PKS’s are usually embedded within their large polypeptides, whereas those of *trans*-AT PKSs are usually encoded on separate, smaller polypeptides.

Most *trans*-AT assembly lines possess Laterally-INteracting Ketosynthase Sequence (LINKS) motifs that link what would otherwise be homodimeric polypeptides into a higher-order architecture^13^ (Figure 1a). These helical motifs are inserted within the KS flanking subdomain (FSD) that is also present in *cis*-AT assembly lines but often referred to as the KS-AT linker^14^. They associate with one another to zipper synthases such as the *Bacillus subtilis* bacillaene *trans*-AT assembly line into organelle-sized megacomplexes, visualizable within cells by EM^15^. Recent cryo-EM studies of portions of the *Brevibacillus brevis* BGC11 *trans*-AT assembly line revealed parallel filaments comprised of the KS and LINKS motifs^16^. The many similarities between *cis*-AT and *trans*-AT assembly lines beg the question whether *cis*-AT assembly lines also possess higher-order architecture.

**Figure 1.**
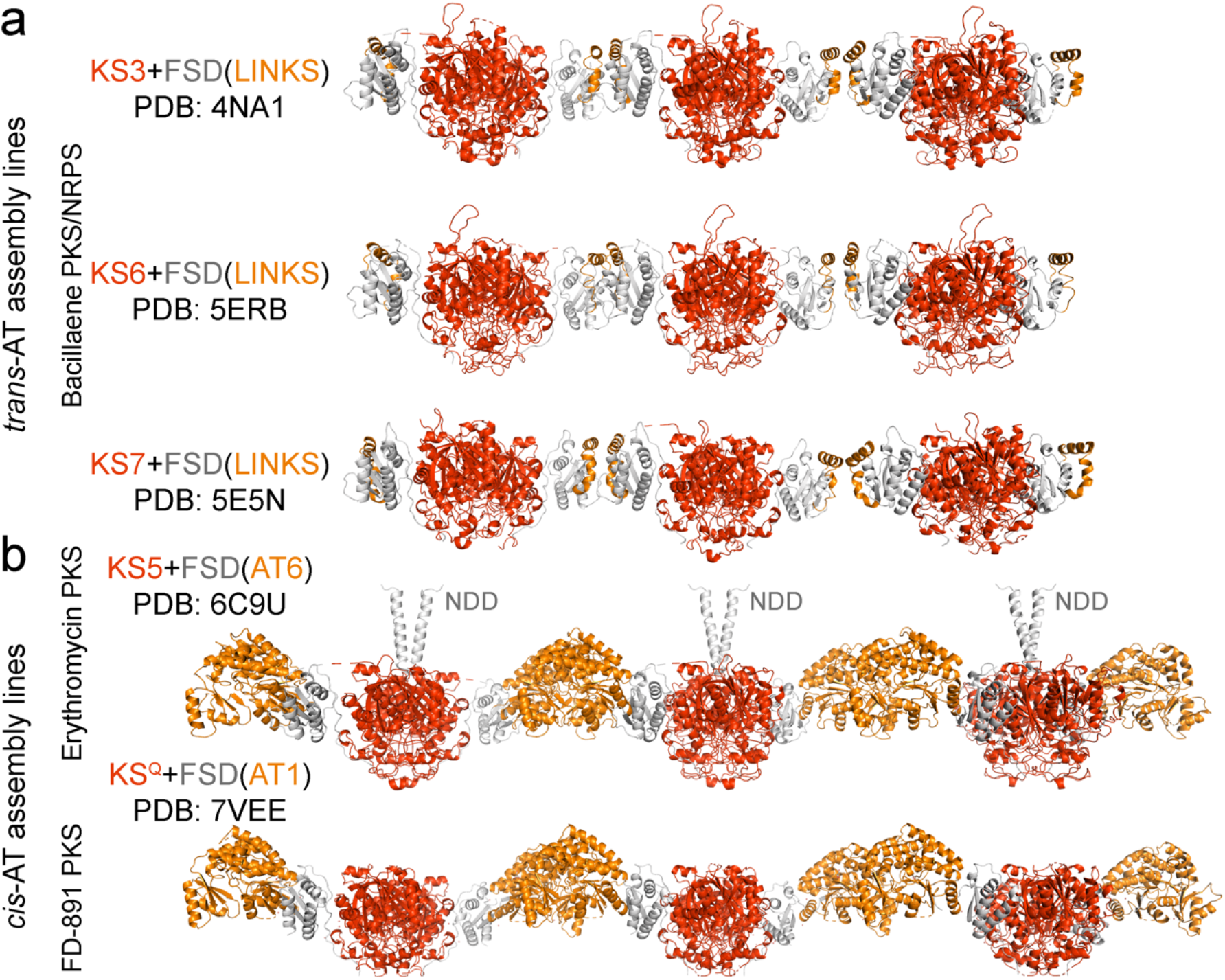
Filaments observed in modular PKS crystal structures. a) Crystal structures of constructs from modules 3, 6, and 7 of related *trans*-AT bacillaene assembly lines show equivalent filaments comprised of KS domains (red) and Laterally-INteracting Ketosynthase Sequence (LINKS) motifs (orange). The LINKS motifs are incorporated within the flanking subdomain (FSD, grey) (PDB: 4NA1, 5ERB, 5E5N). b) Structures from the erythromycin and Gfs *cis*-AT assembly lines (PDB: 6C9U, 7VEE) show equivalent filaments comprised of KS domains (red) and AT domains (orange). The AT domains are incorporated within FSD’s at the same location as LINKS motifs.

Circumstantial evidence indicates that AT domains play a similar role to LINKS motifs to mediate the lateral assembly of *cis*-AT assembly lines. AT domains are inserted within FSD at the same location as LINKS domains, and the angle that AT projects from the KS domain is conserved^5,7,17-19^. Equivalent KS+AT filaments, formed by AT/AT interfaces, appear in three reported crystal structures (PDB: 2QO3, 6C9U, 7VEE)^18,20,21^ (Figure 1b). However, unlike most protein/protein interfaces, few hydrophobic residues are present and little surface area is buried.

Here we present the crystal structure of the most downstream KS+AT didomain of the erythromycin *cis*-AT assembly line at 2.9-Å resolution. Its KS domains are poorly ordered due to the covalent inhibitor cerulenin and do not form contacts with other KS’s as in reported structures, yet its AT domains contact one another as in the filament-containing KS+AT crystal structures. With added confidence that the AT domains of *cis*-AT assembly lines mediate lateral interactions and with assistance from AlphaFold, the framework of the *Streptomyces venezuelae* pikromycin *cis*-AT assembly line was constructed^3,22^. The AT dimers participate in vertical contacts, not only with an upstream KS dimers but also with downstream docking domains^23,24^. The framework creates equivalent, highly-ordered reaction chambers for each module within the synthase. Related crystal structures suggest that the described two-dimensional framework described here represents a layer of a three-dimensional architecture.

## RESULTS & DISCUSSION

### Structure determination

DNA encoding Ery(KS6+AT7) (updated module boundary)^10,11^ was obtained by PCR using pKOS422-31-2^25^ as a template and ligated into a pET28b expression vector. The corresponding histidine-tagged protein was produced by *E. coli* BL21(DE3) and purified by nickel affinity and gel filtration chromatographies. After removal of the histidine tag, Ery(KS6+AT7) was re-purified by gel filtration chromatography. Crystals grew within minutes by hanging-drop vapor diffusion and could not be optimized to diffract beyond 15 Å. Incubating Ery(KS6+AT7) with the covalent inhibitor cerulenin resulted in new conditions and crystals that diffracted to 2.9-Å resolution.

The structure reveals 4 molecules of Ery(KS6+AT7) per asymmetric unit (Figure 2, Table 1). The KS domains are monomeric and possess varying levels of disorder, presumably due to modification of the reactive cysteine, Cys1657, which shifts away from its normal position (Figure 2a). To our knowledge, this is the first report of a KS crystallizing as a monomer. In contrast, each of the AT domains is well-ordered and makes equivalent contact with another AT domain. This contact, observed twice within the asymmetric unit (0.72 rmsd between dimers), is similar to those observed in crystal structures of Ery(KS3+AT4) [PDB: 2QO3 and 6C9U] and Gfs(KS^Q^1+AT1) [PDB: 7VEE]^18,20,21^ (Figure 2b). Compared to these ATs, more side-chains make contact across the dimer interface (Figure 2c)^26^.

**Table 1.** Data collection and refinement statistics.

|  |  |
| --- | --- |
| Wavelength (Å) | 1.116 |
| Resolution range (Å) | 76.23 - 2.9 (3.004 - 2.9) |
| Space group | P 1 21 1 |
| Unit cell (Å, °) | 71.21 167.94 184.23<br>90 99.43 90 |
| Total reflections | 216231 |
| Unique reflections | 93919 (9416) |
| Multiplicity | 2.30 |
| Completeness (%) | 99.24 (99.62) |
| Mean I/sigma(I) | 5.26 |
| Wilson B-factor | 55.2 |
| CC1/2 | 0.836 |
| Reflections used in refinement | 93814 (9382) |
| Reflections used for R-free | 1725 (173) |
| R-work | 0.2440 (0.3494) |
| R-free | 0.3076 (0.4037) |
| Number of non-hydrogen atoms | 22987 |
| macromolecules | 22979 |
| solvent | 8 |
| Protein residues | 3177 |
| RMS(bonds) | 0.01 |
| RMS(angles) | 1.2 |
| Ramachandran favored (%) | 91.77 |
| Ramachandran allowed (%) | 6.5 |
| Ramachandran outliers (%) | 1.73 |
| Rotamer outliers (%) | 0.13 |
| Clashscore | 15.95 |
| Average B-factor | 58.09 |
| macromolecules | 58.09 |
| solvent | 46.56 |

**Figure 2.**
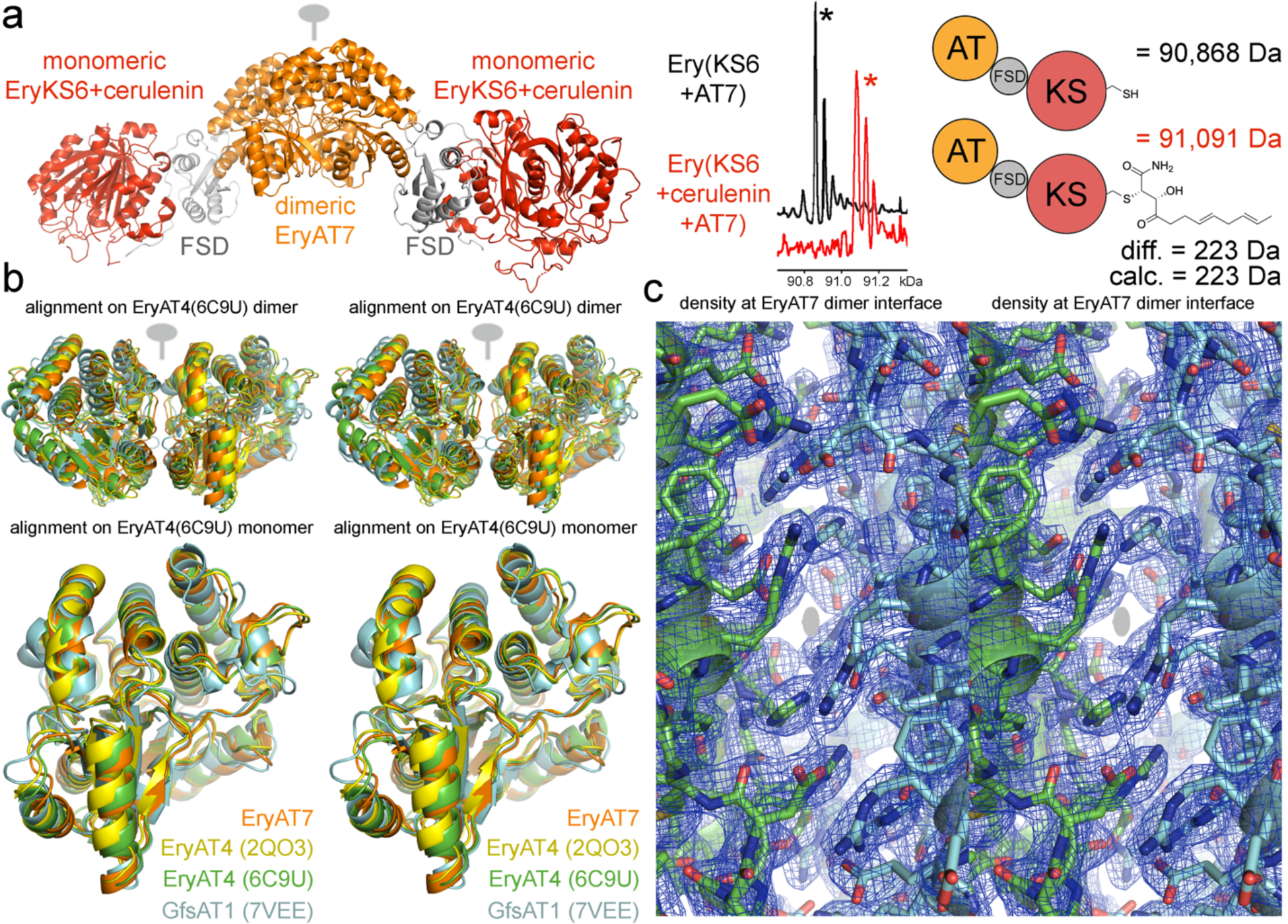
The crystal structure of cerulenin-bound Ery(KS6+AT7). a) The asymmetric unit contains 2 equivalent dimers (0.72 rmsd). The dimer with the lower temperature factors is shown here. Electrospray mass spectrometry shows that Ery(KS6+AT7) was homogeneously modified by the covalent inhibitor cerulenin. b) Few differences between AT dimer interfaces were observed when all known AT dimers were superposed on the EryAT4(6C9U) dimer. Equivalently, when these AT monomers are superposed on EryAT4(6C9U), no major differences are observed. c) Density at the EryAT7/EryAT7 interface, viewed along the twofold axis, reveals it is dominated by charged residues that interact ionically and by stacking. The EryAT7 interface possesses many more contacts than the interfaces observed for EryAT4 or GfsAT1.

### Modeling the framework of the pikromycin PKS

The AT interface in the Ery(KS6+AT7) crystal structure prompted us to model the higher-order architecture of a *cis*-AT assembly line. Additional motivation was provided by recent advances in protein folding prediction^3^. Since our lab is currently performing structural and functional studies with the pikromycin PKS, its architecture was targeted first^7,27,28^. Our study on the loops of *cis*-AT assembly lines revealed that in a PKS homodimer with all of its KS dimers on the same twofold axis the linkers connecting ACP’s and KS’s are not long enough for ACP’s to access their cognate AT’s^29^. However, in a PKS multimer each polypeptide could zigzag to enable ACP’s access to their cognate AT’s (Figure 3). The reaction chambers^30^ would be most rigid and ACP’s would enjoy the most freedom if AT dimers made vertical contacts both with upstream KS dimers and downstream docking domains^23,24^.

**Figure 3.**
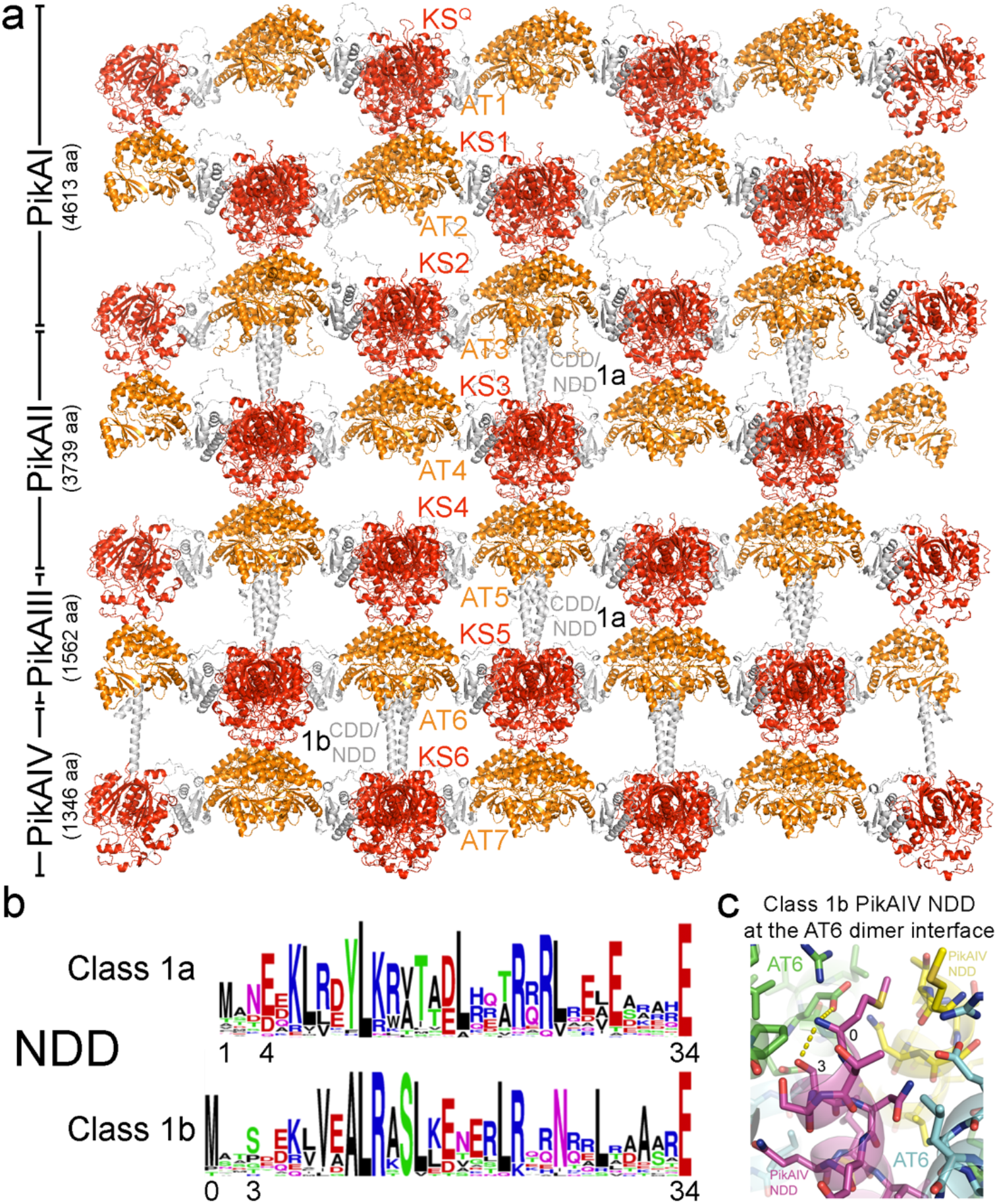
KS, AT, and docking domains form the pikromycin PKS framework. a) Assuming that that the AT/AT interface and KS+AT filaments are relevant to the higher-order architecture of *cis*-AT assembly lines, Ery(KS3+AT4) filaments from PDB 6C9U were used to construct a model. By shifting these filaments half a module width down the assembly line, contact was made between upstream KS dimers and downstream AT dimers. When the domains were replaced with AlphaFold models of the KS, AT, and docking domains of the pikromycin PKS, the NDD/CDD complexes made excellent contact with upstream AT dimers. PDB coordinates for the framework are available (Data Files 1-3). b) Sequence logos of Class 1a and 1b NDD’s reveal the most common location of the N-terminal methionine, 33 residues upstream of the KS domain (the most downstream glutamate is the first KS residue) for Class 1a NDD’s and 34 residues upstream of the KS domain for Class 1b NDD’s. c) The interaction observed at the interface between PikAT6 and the Class 1b NDD of PikAIV suggests that the amine of the initiating methionine interacts with both a conserved serine 4 residues downstream and a conserved glutamate in AT.

To reveal the high-resolution details of the pikromycin scaffold, we constructed an atomic model (Figure 3a). First, a KS+AT filament composed of 6 KS+AT monomers was obtained from the crystal structure of Ery(KS3+AT4) (PDB 6C9U) and oriented so its twofold axes are parallel to the z-axis, with the C-termini facing down^20^. The vector that translates one KS+AT dimer onto a neighboring dimer (length: 145 Å) was determined. A duplicate KS+AT filament was then translated half this vector and down the z-axis until its AT dimers and upstream KS dimers made contact (67 Å). Three copies of the first filament were translated down the z-axis by 134, 268, and 402 Å, and 2 copies of the second filament were translated down the z-axis by 134 and 268 Å. Because AlphaFold generates a conformation slightly different from that observed for KS+AT didomains within filaments, the KS+AT dimers it predicted were divided into KS+FSD dimers and AT monomers. The pikromycin KS+FSD dimers were superposed on appropriate EryKS3 dimers, and the pikromycin AT monomers were superposed on appropriate EryAT4 monomers.

The next step in constructing the atomic model of the pikromycin PKS scaffold was incorporating docking domain complexes comprised of the final residues of C-terminal docking domain (CDD) motifs and the N-terminal docking domain (NDD) motifs^23,24^. By superposing AlphaFold structures of NDD+KS dimers on KS dimers in the model, the NDD coiled coils were positioned. From CDD/NDD complexes generated by AlphaFold, NDD’s and the final residues of CDD’s were then superposed on the positioned NDD coiled coils. Sequence logos^31^ of the Class 1a and 1b NDD’s reveal the most common positions for the N-terminal methionine to be 33 and 34 residues upstream of the KS domain, respectively (Figure 3b). The interface between PikAT6 and the Class 1b NDD of PikAIV shows the methionine amine interacting with both a conserved serine 4 residues downstream and a conserved glutamate in AT(Figure 3c). The domain/domain interfaces in the model were energy-minimized with CHARMM using ClusPro 2.0^32,33^. A homodimeric unit was then selected from the model such that the scaffold could be readily constructed through its translation (coordinates available in Data Files 1-3).

### Interfaces

The pikromycin scaffold contains 3 uncharacterized AT interfaces: the vertical AT/KS interface, the lateral AT/AT interface, and the vertical AT/docking domain interface. To understand them better, those in the pikromycin framework were analyzed with the help of bioinformatics, and the representative PikAT3 interfaces are described here (Figure 4). In case these interfaces vary between PKS’s expressed by different organisms, only sequences from characterized streptomyces PKS’s were employed^9^. Analysis of the first 2 interfaces was aided by sequence logos constructed using 186 γ-modules not split by docking domains from 63 synthases^29,31^. Analysis of the third was aided by sequence logos constructed using 91 γ-modules split by Class 1a docking domains from 64 synthases^29^.

**Figure 4.**
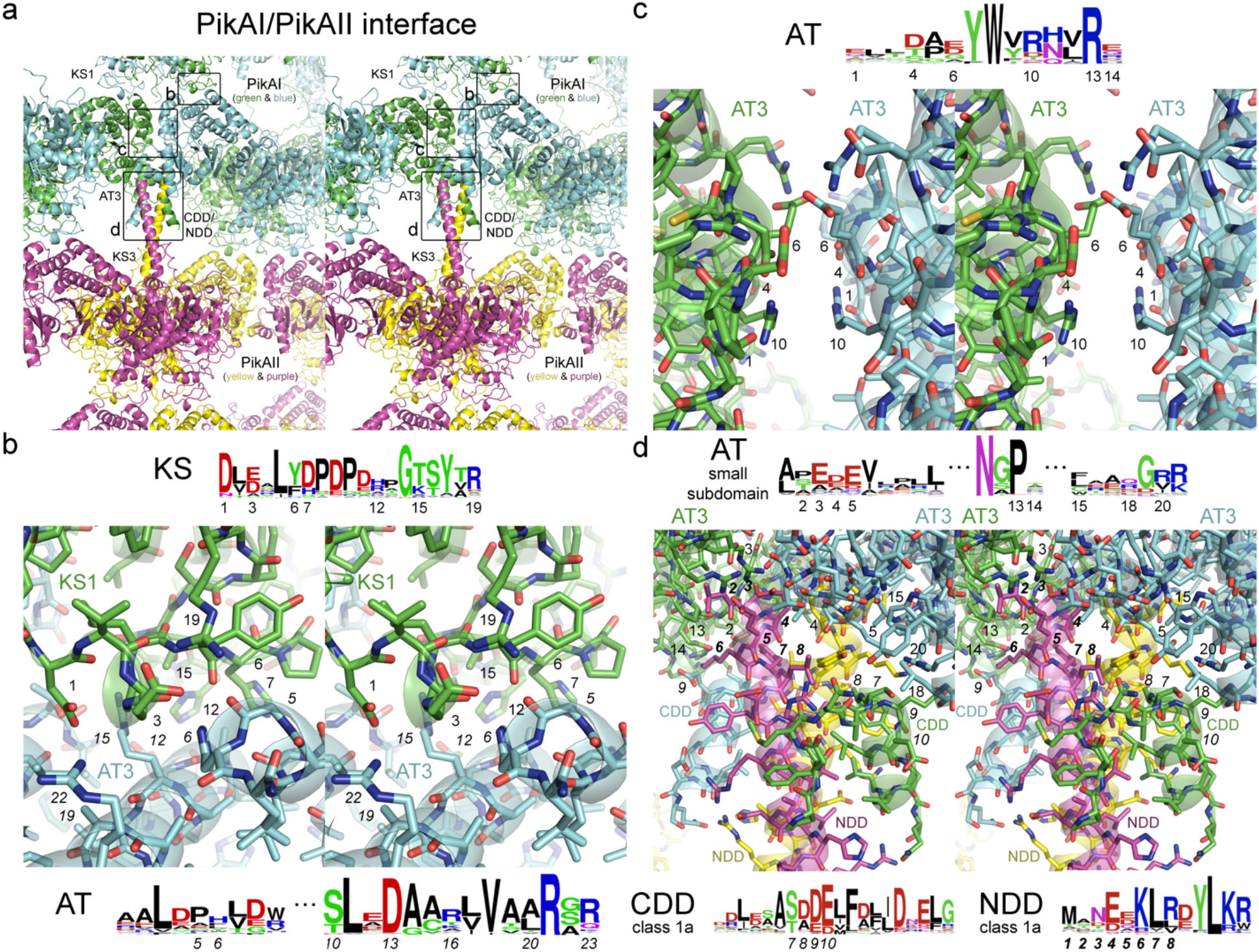
AT interactions at the PikAI/PikAII interface. a) PikAT3 contacts PikKS1, another copy of itself, and the complex between PikAI CDD and PikAII NDD. Letters indicate the panels illustrating the molecular details of the boxed interfaces. b) The interface between PikKS1 and PikAT3 is primarily between KS residues 1151-1169 (positions 1-19, regular font) and large AT subdomain residues 3213-3220 and 3287-3300 (positions *1*-*22*, italic font). Sequence logos show the conservation of key residues, such as the aspartate at position 1 and the arginine at position 19. c) The PikAT3/PikAT3 interface is primarily between large AT subdomain residues 3419-3432 (positions 1-14, regular font). Sequence logos show the conservation of key residues, such as the arginine at position 10 that usually interacts homotypically through the twofold axis of the AT dimer. d) The interface between PikAT3 and the PikAI CDD/PikAII NDD complex is primarily between small AT subdomain residues 3316-3325, 3341-3344, and 3362-3368 (positions 1-21, regular font), PikAI CDD residues 4592-4611 (positions *1*-*20*, italic font), and PikAII NDD residues 3-15 (positions ***1***-***13***, bold italic font). Sequence logos show the conservation of key residues, such as the glutamate at position 3 that may interact with the terminal NDD amino group or the glutamate/arginine pair (positions 5/20) that often helps form the binding pocket for CDD. The 3 N-terminal residues of PikAII could not be confidently modeled; however, a methionine most commonly is at position ***1*** where its amino group can interact with the conserved glutamates at position ***4*** of NDD and position 3 of AT.

#### Vertical interactions between AT’s and KS’s

The pikromycin scaffold contains 5 vertical interfaces between AT and KS domains as well as 1 between an AT and a KS-like domain, KS^Q^, that generates a propionyl primer unit from a methylmalonyl building block^7^ (Figure 3a). Here, the interaction between PikKS1 and PikAT3 helps describe the features of the vertical KS/AT interface. A structured loop formed by residues 1151-1169 (positions 1-19, regular font) on the PikKS1 surface fits into a groove formed by three α-helices on the AT surface (Figure 4b). The KS loop primarily makes contact with residues 3213-3220 and 3287-3300 (positions *1*-*22*, italic font). The conserved aspartate at position 1 of PikKS1 interacts ionically with the conserved arginine at position *22* (in PikKS2-PikKS5 it interacts ionically with the conserved arginine at position *15*). A negatively-charged residue at position 3 helps orient the conserved arginine at position 19 to interact with the carbonyl of position *5*. The residues at positions 6, 7, and 8 make van der Waals interactions with positions *1, 2*, and *5*, and the residue at position 12 makes van der Waals interactions with positions *11* and *12*. The carbonyl of position 13 interacts with a conserved arginine at position *15*. Because residues 1-19 are not conserved in KS^Q^ domains^7^ and the modeled interface between PikKS^Q^ and PikAT2 is the least complementary within the pikromycin synthase, KS^Q^’s and AT1’s may not be rigidly connected to the framework.

#### Lateral interaction between AT’s

As with the AT/AT interfaces observed for EryAT4 and GfsAT1, those of the pikromycin synthase bury little surface area (Figure 4c)^18,20,21^. Most of the contacts are made by residues 3419-3432 (positions 1-14) of the large subdomain^26^. The main contacts are homotypic through the twofold axis of the AT dimer between the side chains of a conserved arginine at position 10^34^ and the residue at position 6. In PikAT3 the arginine is oriented by aspartates at positions 1 and 4; however, in the other pikromycin AT’s it is oriented by a glutamate 16 residues upstream of the arginine. This glutamate also interacts with a histidine at position 14 of the other monomer.

The AT/AT interface observed for EryAT7 buries more surface area than the interfaces observed for EryAT4 and GfsAT1 (Figure 2c). Residues from the small subdomain participate by making contact with residues from the large subdomain of the other monomer (R2156 and E2163 with E2244, R2245, D2276). Contact between the large subdomains also differs from the other observed AT/AT interfaces. While D2258 substitutes for the conserved arginine at position 10, R2254 at position 6 makes both homotypic contact through the twofold and an ionic interaction with D2250 at position 2 of the other monomer. The interface also features 2 equivalent stacks of side chains, comprised of L2241, H2242, R2246 from one monomer and R2261 (position 13) from the other. Several smaller contacts are also made by ionic and polar side chains.

#### Vertical interaction between AT’s and docking domains

The unusual domain-domain interface observed for the AT dimer is understood better when its role as a receptor for CDD/NDD complexes is considered (Figure 4d). The AT dimer must bind to the N-terminal end of the NDD coiled coil. The most common lengths of NDD’s are 33 and 34 residues for Class 1a and 1b NDD’s, respectively (Figures 3b and 3c). The residue at the fourth position minimizes fraying of the NDD by interacting with the amino group of the initiating methionine. This preorganization would help the binding of the docking domain complex and the AT dimer. NDD’s are frequently longer than 33-34 residues (Class 1a moreso than Class 1b) and the AT dimer interface must accommodate the additional residues.

The interface between PikAT3 and the PikAI CDD/PikAII NDD complex is primarily between small AT subdomain residues 3316-3325, 3341-3344, and 3362-3368 (positions 1-21, regular font), PikAI CDD residues 4592-4611 (positions *1*-*20*, italic font), and PikAII NDD residues 3-15 (positions ***1***-***13***, bold italic font). While the first 3 residues of PikAII could not be confidently modeled, a methionine at NDD position ***1*** usually interacts with a conserved glutamate at position ***4*** of NDD and a conserved glutamate at position 3 of AT, oriented by a conserved AT arginine. The side chain of NDD position ***2*** makes van der Waals contact with the conserved proline at position 13 of AT. The asparagine at NDD position ***3*** interacts with the side chain at position 2 of AT. The side chain of NDD position 4 makes van der Waals contact with the side chains at position 4 of AT. The glutamate, lysine, and tyrosine of NDD positions 5-7 make subtle interactions with nonconserved AT side chains. CDD residues *7*-*10* fill a small pocket on AT formed by residues 5, 9, 14, 15, 18, and 20. The model of the pikromycin framework suggests this interaction is stronger for Class 1b CDD’s. The residue at CDD position *7* (serine in Class 1a, aspartate in Class 1b) has the dual role of capping the final polypeptide helix and making the principal contact with the AT pocket.

#### Interactions between modules

Polypeptides zigzag along the assembly line such that a module in the middle of a synthase makes contact with 2 modules upstream of it and 2 modules downstream of it (Figure 5). Its interface with the downstream module lies between its KS and the FSD of the downstream module and is primarily made through interactions between KS surface residues and the FSD LPTYxF motif. Its interface with the module 2 modules downstream of it is the interface between KS and AT described here (Figure 4b). Engineering experiments using the updated module boundary have shown the modularity of both interfaces^27,28^.

**Figure 5.**
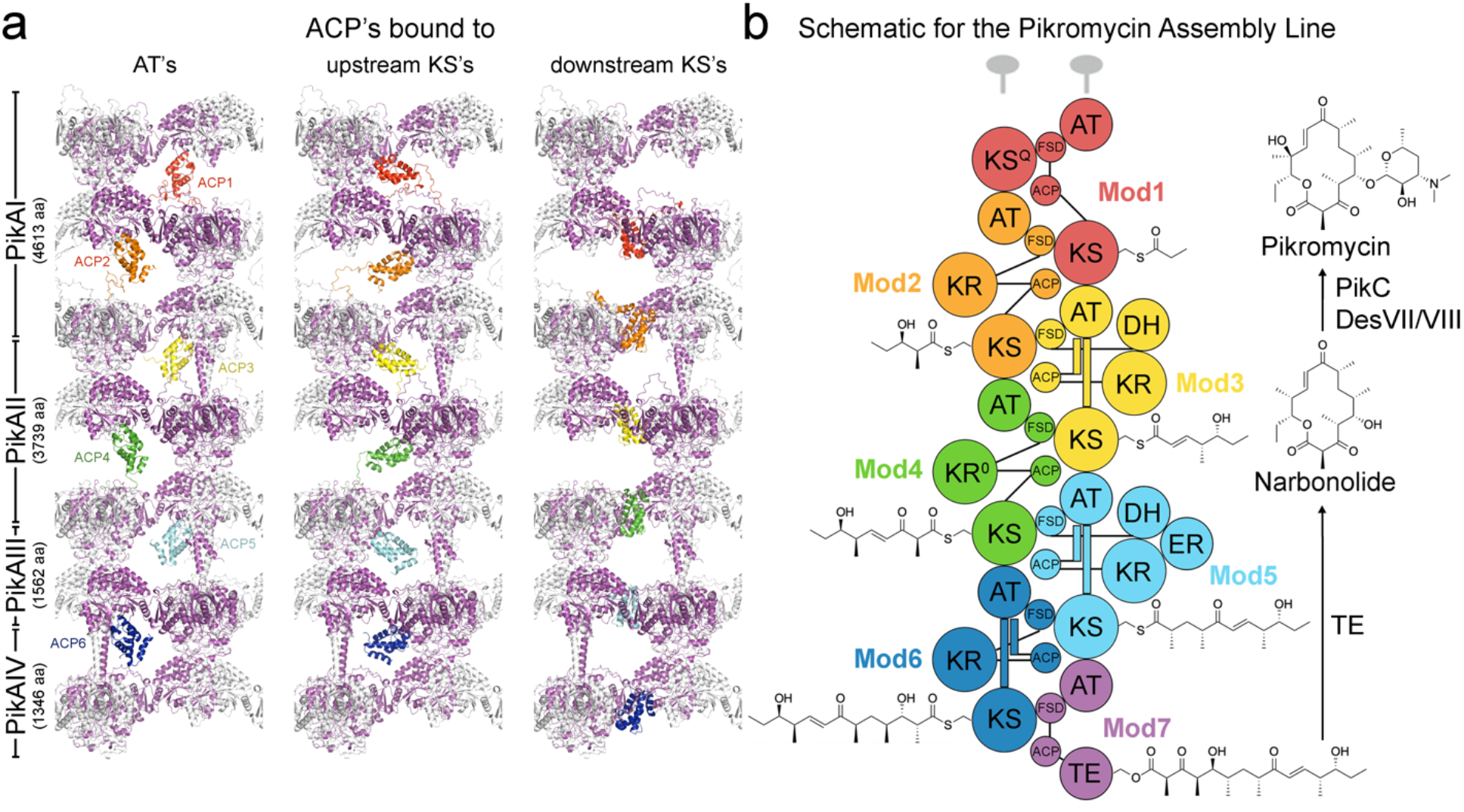
Visualizing ACP movement within *cis*-AT assembly line reaction chambers. a) From left to right, PikACP1-PikACP6 (red-blue) are bound to their cognate AT’s to collect malonyl and methylmalonyl extender units, upstream KS’s to fuse these extender units with growing polyketide chains, and downstream KS’s to verify processing of the extended chain and pass it to the downstream module through a subsequent chain extension reaction. Only a portion of the synthase is shown here since its 6 reaction chambers are equivalent throughout the synthase. The purple polypeptides indicate PikAI-PikAIV monomers. While AT acts in *cis*, the upstream KS and the downstream KS act in *trans*. ACP interfaces with the AT and the downstream KS are predicted by AlphaFold, whereas the interface with the upstream KS has been observed by cryo-EM^5,6^. ACP linkers were built for those anchored on the framework (*e*.*g*., the linkers for PikACP1 connect to C-terminal end of PikFSD1 and the N-terminal end of PikKS1). A movie showing how ACP’s move to construct polyketides in a hand-over-hand manner was generated (Movie S1). b) A schematic of PikAI-PikAIV monomers, colored using the updated module boundary, indicates the reaction chambers of the synthase as well as how the PKS framework is constructed by applying the twofold axes. The NDD’s and CDD’s are indicated with long and short helices, respectively. The location of processing enzymes remains mysterious. ACP linkers are indicated with black lines.

### Adding ACP’s and linkers to the framework

To begin understanding how ACP’s move within their reaction chambers, 6 pikromycin ACP’s and 4 of their linkers were added to the framework (Figure 5). The ACP’s were generated by AlphaFold and placed at the experimentally-determined extension docking site on the upstream KS^5,6^ as well as at the AlphaFold-predicted docking sites for AT and the downstream KS (Bista et al., manuscript in preparation; Hirsch et al., manuscript submitted). Since only ACP linkers anchored on the framework were added, only PikACP1 is shown with both its N- and C-terminal linkers. All of the modeled linkers still possess ample slack with the ACP’s bound to their cognate upstream KS’s, AT’s, and downstream KS’s. A movie showing the hand-over-hand synthesis of the pikromycin precursor, narbonolide, was generated (Movie S1).

### The third dimension

The framework presented here for the pikromycin synthase does not include any of its processing enzymes, the first portion of its CDD’s, or its thioesterase (TE). DH’s, the first portion of CDD, and most TE’s are dimeric, and most synthases also possess dimerization elements (DE’s) upstream of β-module KR’s^1,35^. With the probable exception of TE, these dimeric features do not fit on the twofold axes of the described framework. Thus, the framework presented here may be just one layer of the higher-order architecture of *cis*-AT PKS’s, with DH’s, DE’s, and the first portion of CDD’s connecting the layers through their dimer interfaces. The separation between the layers must be wide enough for *trans*-enzymes to access polyketide intermediates^9^.

Some synthases may have the ability to grow infinitely in three dimensions. Pks12 from *Mycobacterium tuberculosis* is a 1-polypeptide, 2-module synthase that performs 10 extensions in the synthesis of mycoketides^36^. Since it possesses an NDD and a CDD, it is thought to multimerize along its vertical axis. Its AT’s, DH’s, and the first portion of its CDD may enable it to multimerize in both lateral dimensions as well. Atomic force microscopy analysis of this synthase indicates that it possesses higher-order organization.

## Conclusions

The described higher-order architecture for *cis*-AT assembly lines continues the trend started by the α_6_β_6_ fungal FAS of higher oligomerization states enabling the formation of equivalent, well-organized reaction chambers^30^. As both *cis*-AT and *trans*-AT assembly lines frequently contain nonribosomal peptide synthetase (NRPS) modules, it seems likely that NRPS assembly lines also possess higher-order architecture and well-organized reaction chambers. Equivalent dimeric NRPS structures have been observed^37,38^. It is also conceivable that iterative PKS’s, including the mammalian FAS, multimerize to form well-organized reaction chambers^39^.

A better understanding of *cis*-AT PKS architecture will empower PKS engineering. By structurally modeling desired hybrid synthases before genetically constructing them, incompatibilities may be detected and solutions to overcome them identified. It is through connecting the functions of these molecule factories with their structures, as complex as they may be, that the promise of harnessing them to generate new molecules and medicines will be realized.

## METHODS

### Cloning, expression, purification, and crystallography

The expression plasmid for N-terminally histidine-tagged Ery(KS6+AT7) was constructed by cloning the corresponding DNA from pKOS422-31-2 using primers 5’-GGAGATATACATATGGCCGATGATCCAATTGCGATCGTAGGTATGGCCTGTC-3’ and 5’-GTGGTGCTCGAGTCATCACACCTCCGGCTGCAGCCAG-3’, digesting it with NdeI and XhoI, and ligating it between those sites in pET28b. After transforming *E. coli* BL21(DE3) with it, cells were grown at 37 °C to OD_600_=0.4, induced with 1 mM IPTG, and left 14 h at 15 °C. They were then collected and sonicated in lysis buffer (500 mM NaCl, 30 mM Tris pH 7.4), and the lysate was poured over a nickel-NTA column equilibrated with lysis buffer. After washing with 15 mM imidazole in lysis buffer, the protein was eluted with 150 mM imidazole in lysis buffer. After concentration, the protein was injected onto a Superdex 200 gel filtration column equilibrated with 150 mM NaCl, 10 mM Tris pH 7.4. The histidine tag was cleaved with thrombin, which was subsequently removed with a benzamidine-sepharose column. After passing the protein through the Superdex 200 column again, it was collected and concentrated to 8 mg/mL.

Crystals were obtained in 22% PEG20K, 100 mM Tris pH 8.0 but did not diffract beyond 15 Å even after standard optimization techniques. These crystals could be observed to grow within minutes. New crystallization hits were obtained in the presence of cerulenin. Electrospray mass spectrometry confirmed the homogeneous modification of Ery(KS6+AT7) (Figure 2a). Ery(KS6+AT7) was supplied with 1 mM cerulenin and added 3:1 to the well condition (10% PEG3350, 150 mM MgSO_4_, 150 mM NaCl, 100 mM Tris pH 7.5) in a hanging drop vapor diffusion format at 22 °C. Clusters of thin plates grew to full size over one month. Surgery was necessary to obtain single crystals suitable for mounting. Crystals were cryoprotected in 20% glycerol, 15% PEG 3350, 150 mM MgSO_4_, 150 mM NaCl, 100 mM Tris pH 7.5 prior to flash-freezing in liquid nitrogen. Data were obtained at ALS Beamline 8.3.1.

Images were processed with XDS^40^. The structure was solved by molecular replacement, searching for 4 KS monomers and 4 FSD+AT monomers obtained from the structure of Ery(KS5+AT6) (PDB 2HG4)^17^ using PHENIX^41^. Modeling and refinement were performed with Coot^42^ and PHENIX. An AlphaFold model^3^ of Ery(KS6+AT7) helped construct regions with weak density (Table 1).

### Modeling the pikromycin PKS framework and ACP’s

The framework was constructed as described using a combination of PyMOL (Schrödinger, LLC), MOLEMAN^43^, and AlphaFold^3^. CHARMM energy-minimization of the interfaces was performed by ClusPro^33^. ACP’s and their linkers were added using a combination of AlphaFold and Coot^42^.

## ACKNOWLEDGMENTS

This work was supported by the NIH (GM106112) and the Welch Foundation (F-1712). James M. Holton helped process the crystallographic data.

## AUTHOR CONTRIBUTIONS

A.T.K. cloned, purified, and solved the structure of Ery(KS6+AT7). A.T.K., T.M., J.Z., K.A.R., J.L., and R.M. constructed the model of the pikromycin framework. A.T.K. wrote the manuscript. S.N.L. and A.T.K. generated Movie S1.

## DATA AVAILABILITY

Coordinates and structure factors were deposited under PDB code 8G4U.

## SUPPLEMENTARY MATERIAL

The framework coordinates (Data Files 1-3) and polyketide biosynthesis movie (Movie S1) are publicly available.

